# In Vivo Assessments of Minocycline combined Hyaluronic Acid-mediated Ultrasound Therapy of Infected Wound in Rats

**DOI:** 10.1101/450585

**Authors:** Hongbo Zhang, Deshu Zhuang

## Abstract

**Objectives:** The purpose of our research was to examine the effects of Minocycline combined hyaluronic acid (HA)-mediated Ultrasound therapy of infected wound in wister rats.

**Methods:** 40 female wister rats were made wound on the two side of the backbone, then infected in Staphylococcus aureus at the comic for three times. then, they are divided into four groups: control group, minocycline combined HA alone, ultasound alone, minocycline combined HA-mediated ultasound group, respective. After 3 times of treatments, the rats were killed and made into specimens. Assessments consisted of visual inspection in the change of the skin, scar formation pathological morphology by hematoxylin and eosin(HE) stain with optical microscopy, IL-1B assaying and TNF-a were performed.

**Result:** Compared with control group, minocycline combined HA alone, ultasound alone, minocycline combined HA-mediated ultasound group all have effect for wound healing, there was a obvious improvement in all parameters over the duration of the experiment(P<0.05). Compared with the control group, minocycline combined HA-mediated ultasound group indicated less inflammation cells (P<0.001) and the reduce of and IL-1B and TNF-a (P<0.001).

**Conclusion:** Minocycline combined HA-mediated ultrasound can accelerate tissue regrowth, which exert significant benefits in healing the wounds.

## Introduction

Wound infection is the body tissue damage caused by various injury factors which is a common dynamic process that begins from the moment of injury, lack of specificity and useful treatment [1–2]. when the body is suffered wound infection, a series of complex cell biology effects has been started in one’s body, including activation and transfer of stress signals, imbalance of inflammations, release of various harmful factors and etc.inflammatory cytokines are sensitive indicators of post-traumatic inflammation, which includes TNFA, IL-1, IL-6, C-reactive protein and etc[3–5]. By monitoring of these important indicators, we can evaluate the severity of the inflammation, and know the development of inflammatory. By far, the most common treatment for wound infection is surgical debridement, which can eliminate the inflammation rigorously but has avoidless cut, and has high recurrence rate[6].

Hyaluronic acid (HA) is an important component of extracellular matrix. thanks to a number of favourable properties: virtually no toxicity, lack of immunogenicity, anti-inflammatory character, biodegradability, and at the same time provision of high osmotic pressure and hydration[7–9]. In the last twenty years, HA has increasingly been studied for targeted therapies, since its most important receptor, CD44, is over-expressed in a number of cancers, and is heavily involved in HA endocytosis[10]. CD44 is also expressed on activated inflammatory cells, in particular on macrophages. Indeed HA-based formulations have already shown efficacy for the targeted and intracellular delivery of antimicrobials[11]. Minocycline is a second-generation, semisynthetic tetracycline that exerts anti-inflammatory effects which are completely separate from its antimicrobial actions. It showed effective antimicrobial activity aginst staphylococcus aureus and other anaerobic bacteria[12–13]. Previous studies in our group show that the combination of two drug improve the antibiotic activity.control the bacteria, at the same time, contribute the healing of the issue,which is a innovation to treat infect wound.

Ultrasound has been used in medicine for long time ago. Ultrasonic equipment is very simple and cheap,it is a noninvasive technology which first used in medical imaging as a kind of diagnostic tool, in recent years, it is used in trauma and different kind of cancers[14–16]. ultrasound can access into the deep-seated tissue, because the effects of sonication are localized to pathological sites, it can minimize the damage to surrounding normal tissues. In recent years, ultrasound has become a preferred clinical technique in the regulation of targeted therapy[17–19].

Although Minocycline combined hyaluronic acid has been shown effective on wound healing, Minocycline combined hyaluronic acid-mediated Ultrasound on wound healing has not been reported to date. the purpose of our study was to perform if Minocycline combined hyaluronic acid-mediated Ultrasound can improve wound healing in rats.we attempt to provide an experimental basis for future clinical application of Minocycline combined hyaluronic acid-mediated Ultrasound in the treatment of wound healing in humans.

## Materials and methods

### Animals

All the experimental procedures were in accordance with the Institutional Animal Care and Use Committee of Harbin Medical University (Harbin, People’s Republic of China). The protocol was approved by the Experimental Animal Ethics Committee of Harbin Medical University. The operation procedures were performed under 10% chloral hydrate anesthesia. Fourty healthy male Wistar rats, weighing at 240g- 250 g, were used in this experiment. All animals were artificially fed according to the regulations of the Home Office of the Chinese Government, and were given enough food and water. Rats were purchased from The Animal Facilities, Fourth Affiliated Hospital of Harbin Medical University.

### Bacteria cultivate

Staphylococcus aureus(ATCC6538) is choosed to be our infectious bacteria. The bacterial suspensions were incubated in Luria-Bertani(LB) broth at 37□. After 24h culture, the bacterial suspensions were diluted with phosphate-buffered saline (PBS) solution to a final concentration of 1010 cells/ml corresponding to an optical density. Build of infection model

General anesthesia was obtained with 10% chloral hydrate (5 mL/kg) via intraperitoneal injection, the infection model was made at the size of 1.2cm*1.2 cm, depth at 0.5 cm, on both sides of the backbone, then infected Staphylococcus aureus at the concentration of 108 for 3 times every other day(Figure 1).

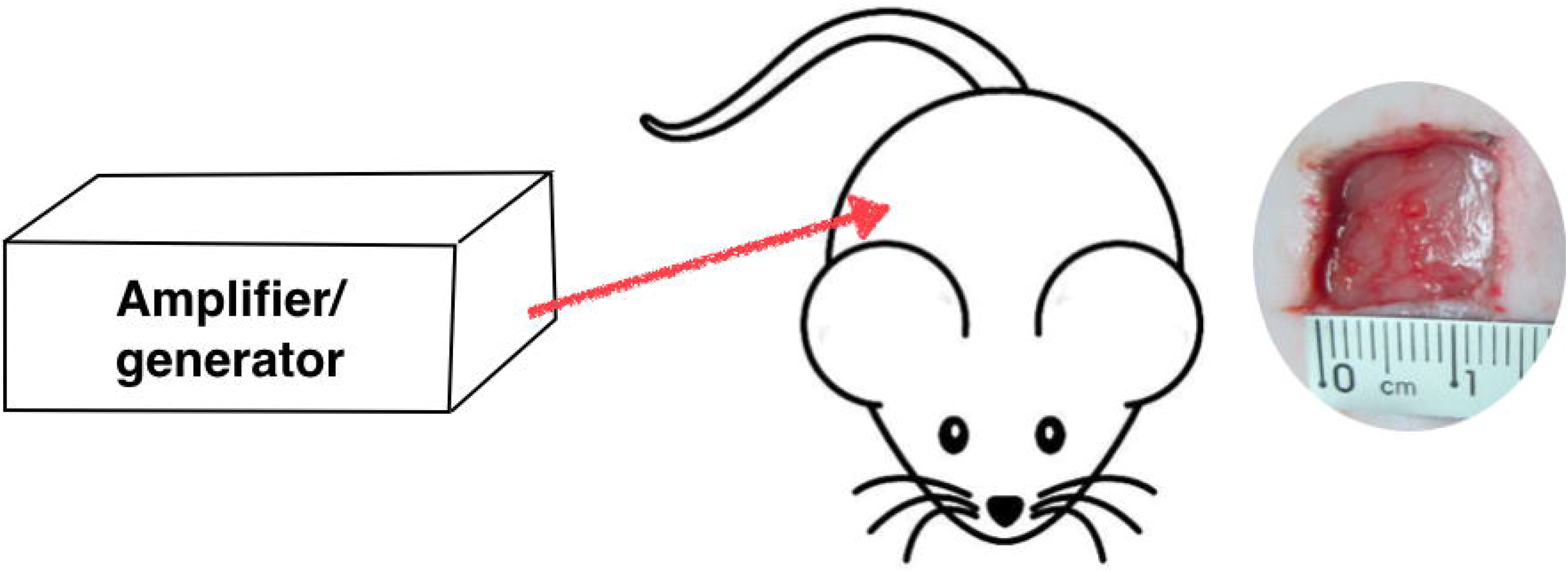

### Ultrasound and medicine

The ultrasound generator and power amplifier used in this study was designed and assembled by Harbin Institute of Technology (Harbin, People’s Republic of China). Ultrasonic intensities (W/cm^2^) were expressed as I_SPTP_ (spatial peak/temporal peak) measured 6 mm away from the ultrasonic transducer radiating surface in the degassed distilled water by a hydrophone (Onda Corp., Sunnyvale, CA, USA). In our experiment, the ultrasonic transducer (diameter: 6 mm; frequency: 1.0 MHz; duty factor: 10%; pulse repetition frequency: 100 Hz) was placed over the wound regions with a medical ultrasonic coupling agent. the ultrasound was placed over the infected wound for 15min with the frequency set at 1W/cm2. Minocycline combined hyaluronic acid gel was delivered as our experiment medicine, with the concentration of 0.1% and 1.5% respectively (Dr Hui Biotechnology Co Ltd, Hangzhou people’s Republic of China).sealed in the −4□ refrigerator.

### Experimental grouping

The rats was randomly divided into four groups(20 samples/group): group 1, no treatment was performed on the wound; group2, Minocycline combined hyaluronic acid on the wound with the concentration of 0.1% and 1.5% respectively; group 3, rats were treated with low-intensity ultrasound therapy (1w/cm2); and group 4, rats with Minocycline combined hyaluronic acid-mediated.

### General observation and determination of wound size reduction

General observations including growth status, color, texture and thickness of the wound healing. there were recorded by taking pictures at designated time points. In addition, after 6 days of treatments, the rats were sacrificed, and the wound sized were measured. the percentage wound size-reduction was calculated using the following formal: Cn =[(S0-Sn)/S0]*100 where Cn is the percentage of wound size-reduction after treatments, S0 is on behalf of the initial wound size, Sn is on behalf of the wound size after treatment.

### Histological and Immunochemical staining

After clinical observation and measure the size of the wound, Normal skin and scar tissue fragments were collected using a scalpel. Skin samples removed for histological analysis were preserved in Karnovsky’s solution (2% paraformaldehyde and 2.5% glutaraldehyde in 0.1 M sodium phosphate buffer, pH 7.2) for 48 h. The samples were dehydrated in ethanol, cleared inxylol, and embedded in paraffin. Cuts of 4-μm thickness were obtained with a rotary microtome (Leica Multi cut 2045_®_; Reichert-Jung Products, Jena, Germany), using one out of every 20 sections to avoid repeating the analysis of the same histological area (Gonçalves et al., 2012, 2014).

For each wound, three serial sections were placed on a slide and stained with hematoxylin and eosin (HE),,and 1L-1P (1 : 75, Leica, England),and TNF-a(). five equidistant sections of each specimen block were selected and captured by a digital camera. Positive and negative immunohistochemistry controls were routinely used. The reproducibility of the staining was confirmed by re-immunostaining via the same method in multiple, randomly selected specimens. The area of wound infection region was histometrically determined using an image analysis system (Image Tool; Harbin Medical University).

### Statistical data analysis

The hypothesis that there has great differences in Minocycline combined hyaluronic acid-mediated Ultrasound therapy of infected wound in rats between other treatment groups was tested using χ^2^ test or Fisher’s exact test. the histometric data were analyzed using the Shapiro–Wilk test, and the intragroup and intergroup analyses were performed with a two-way analysis of variance (*P*< 0.05). When the analysis of variance detected a statistical difference, multiple comparisons were performed using Student’s t test (P <0.05). The derence was considered statistically signigcant if *p* < 0.05. All statistical analyses were performed using SPSS 18 (SPSS Inc., Chicago, IL, USA) software.

## Results

### General observation and wound size reduction

We observed the development of wound infection scars, In our model, wound healing was observed after 7 days of treatment before the sacrifice. in the control group, the scleroma with pyorrhea was obviously raised above the surface of normal skin, and the hyperplasia exceed the scope of the original wound. minocycline combined HA alone, ultasound alone groups, shows better effect compare to the control group, with scleroma vanish, pyorrhea reduce, and with size reduce; while minocycline combined HA-mediated ultasound group scars exhibited obvious size reduce and a gradual softening (Figure 2).

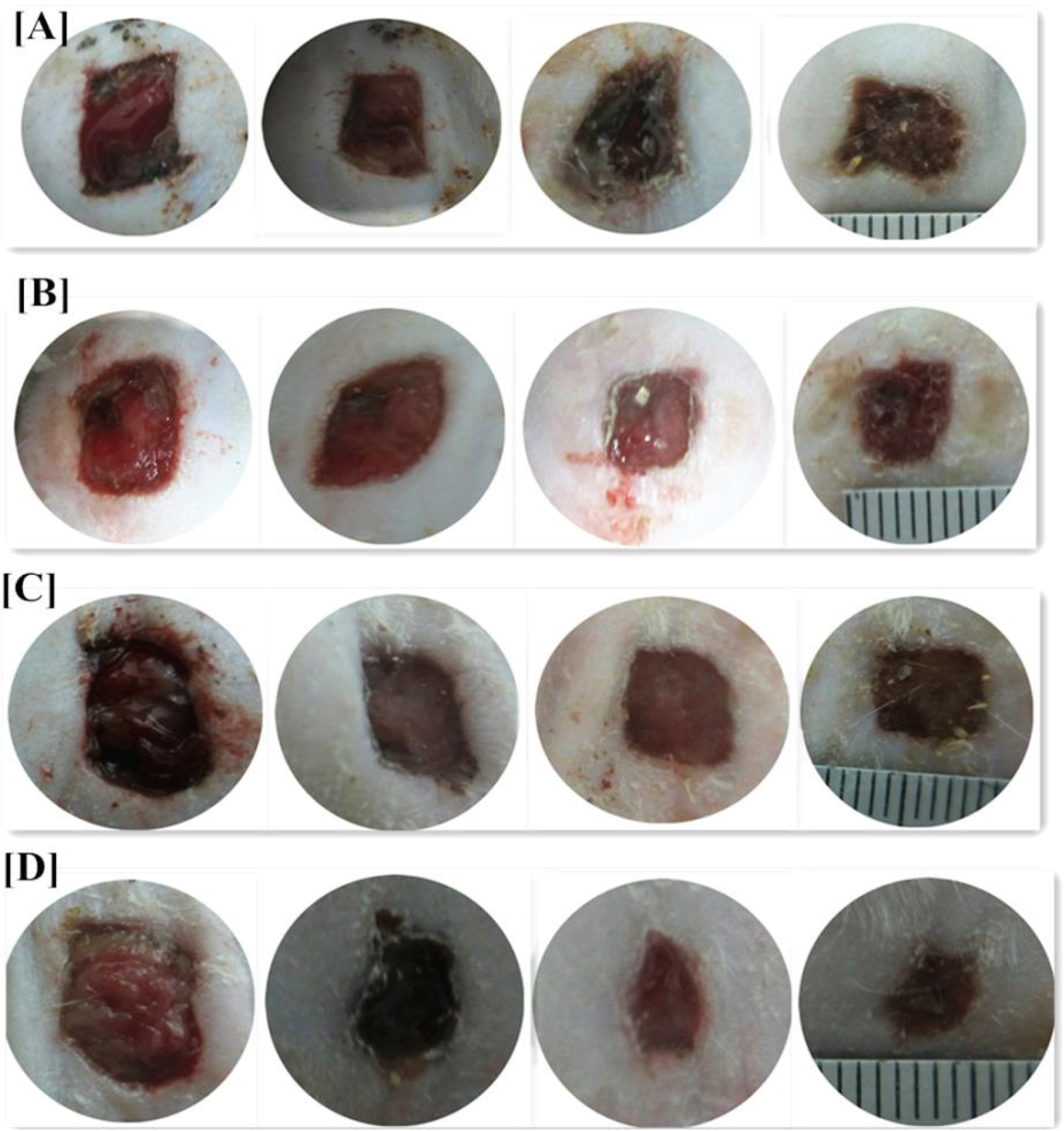

Wound areas of all the treatment groups were smaller than in the control group(Figure 3). minocycline combined HA-mediated ultasound group had best results compared to the others while the percentage of wound size-reduction is 73.89%(P<0.001), In the minocycline combined HA alone group, the percentage of wound size-reduction was sightly bigger than in the ultrasound alone group, but this difference was not statistical significant(P>0.05).

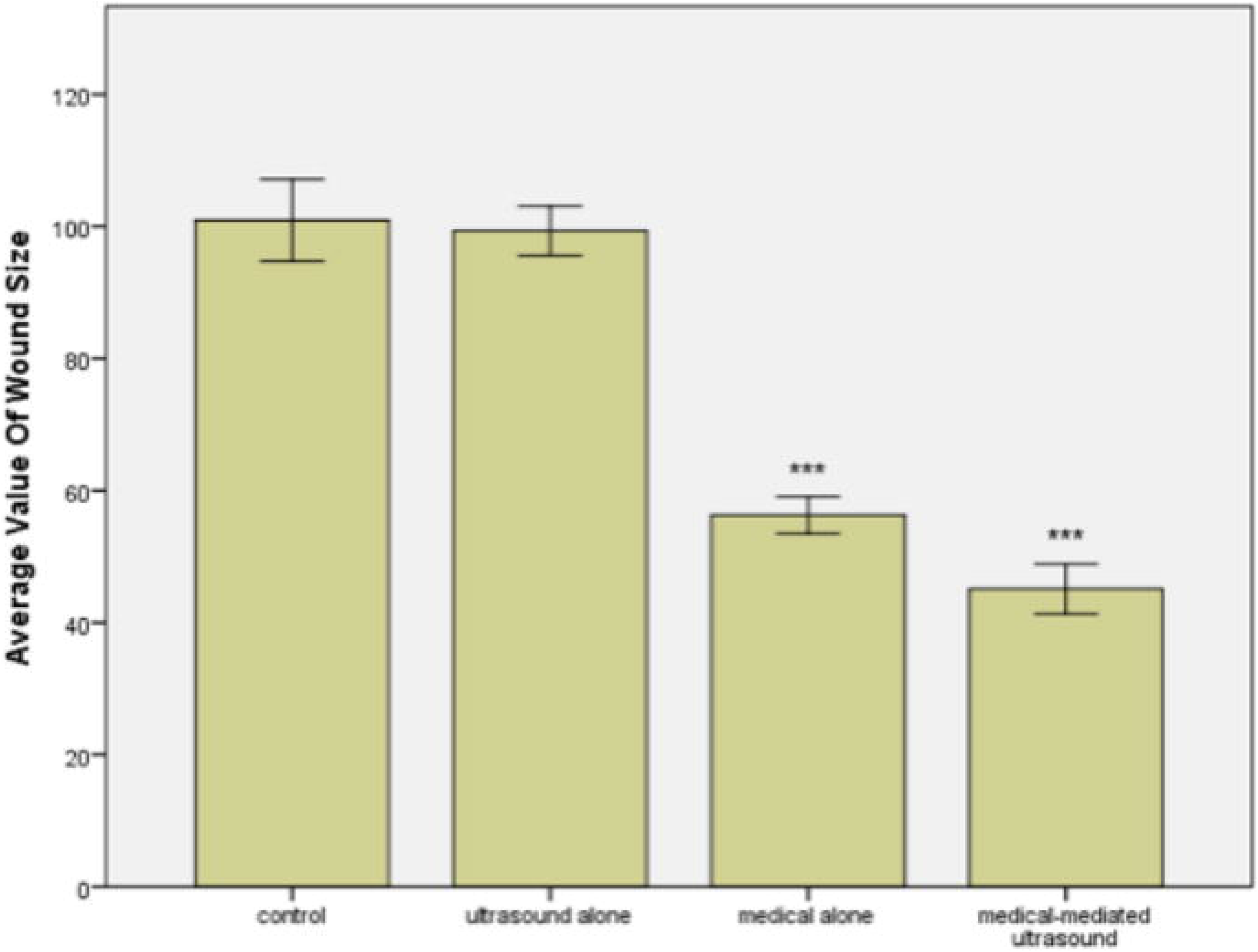

### Histologic examination

The histological appearance of the wound tissue at low magnification in the four groups is shown in Figs. 4. in the control group dermal tissue revealed a large number of inflammatory cells, and disarranged collagen with nodule or vortex-like distribution.in the ultrasound also revealed some inflammatory cells, which mainly in the deep tissue.minocycline combined HA group revealed obvious reduction in the number of inflammatory cells (P<0.001),and a large number of fibroblasts in the deeper tissue.in the minocycline combined HA-mediated ultasound group, revealed obvious reduction in inflammatory cells(p<0.001),ordered arrangement of their collagen fibers, and maturation of granulation tissue.

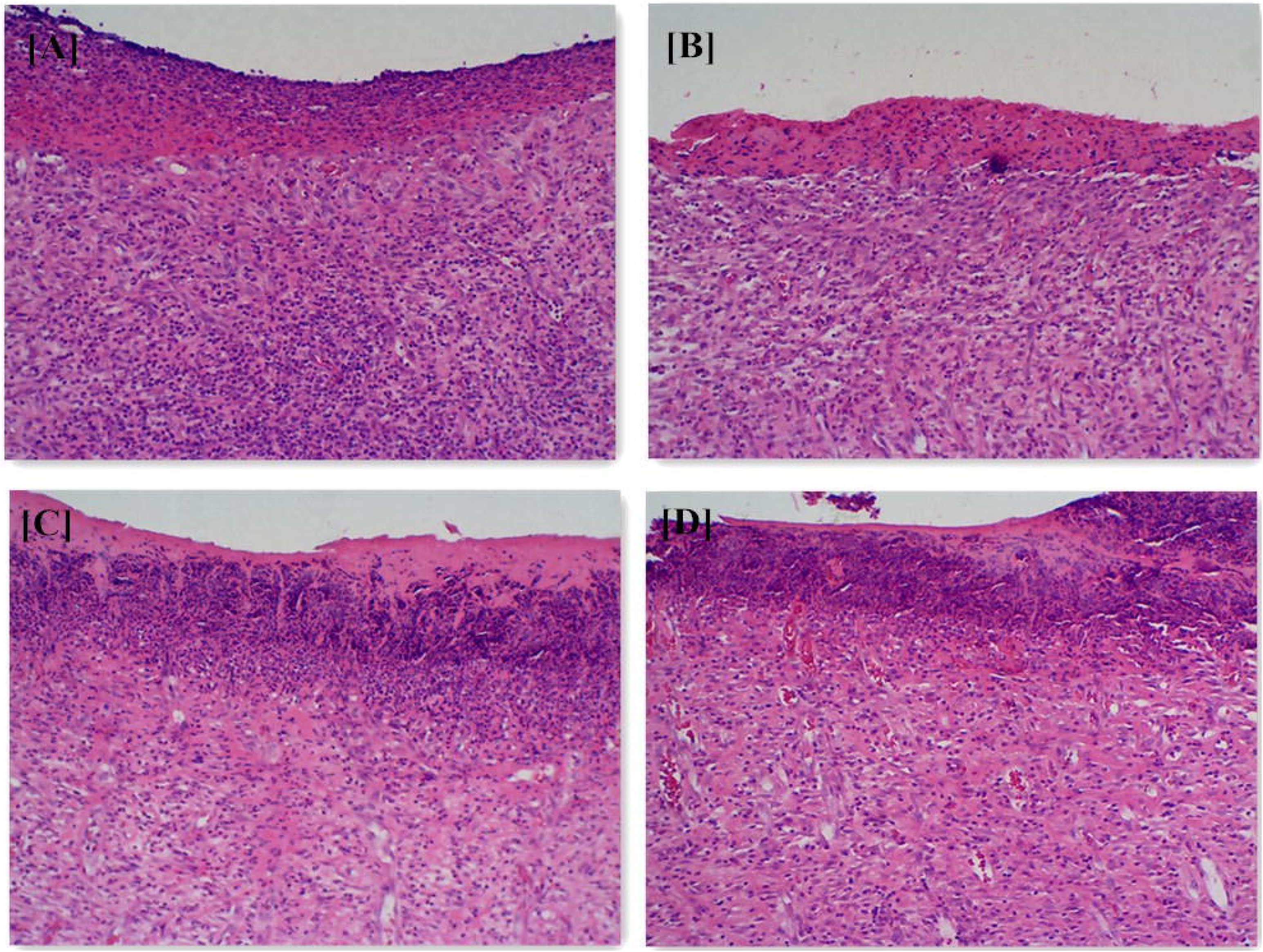

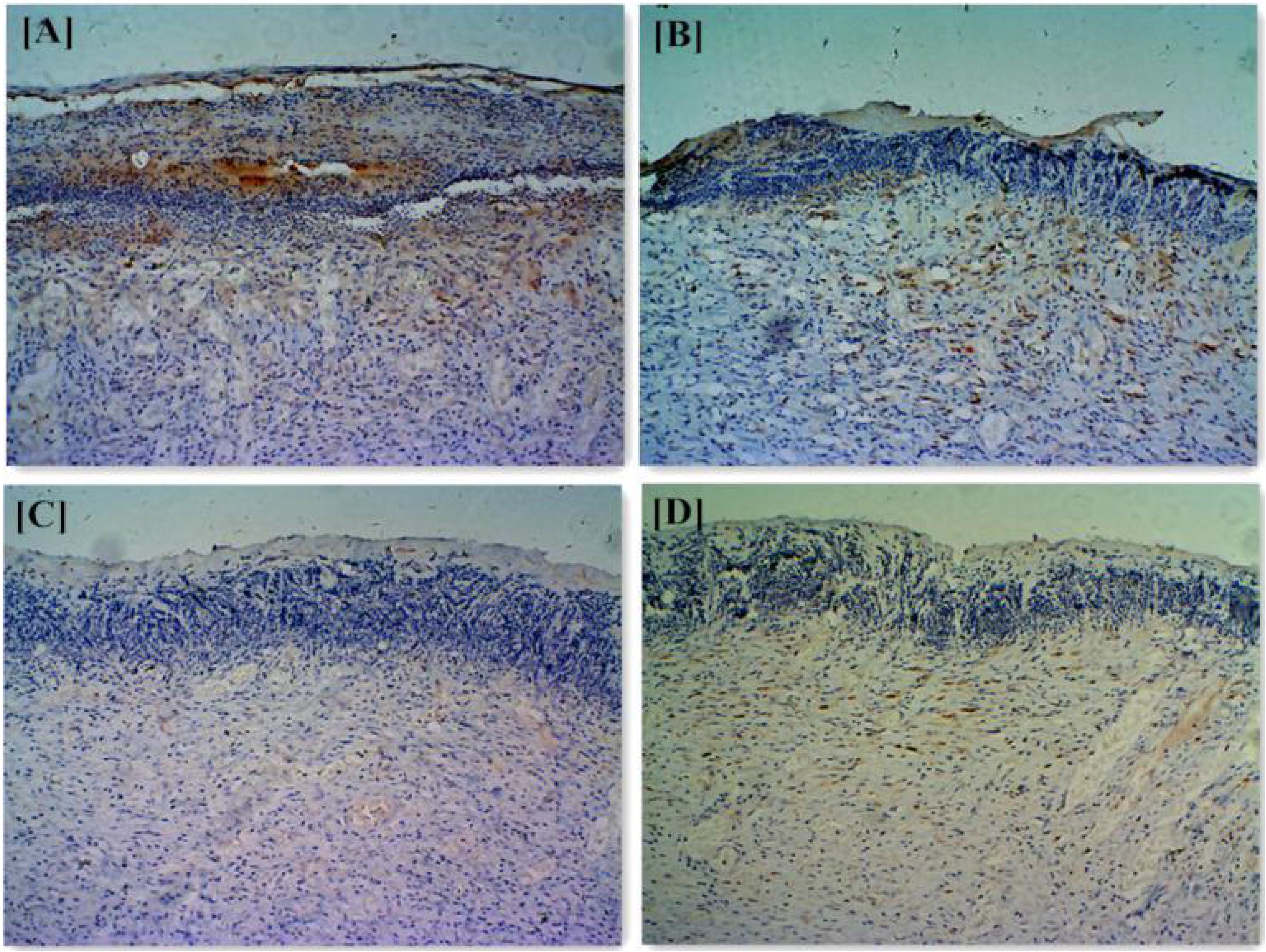

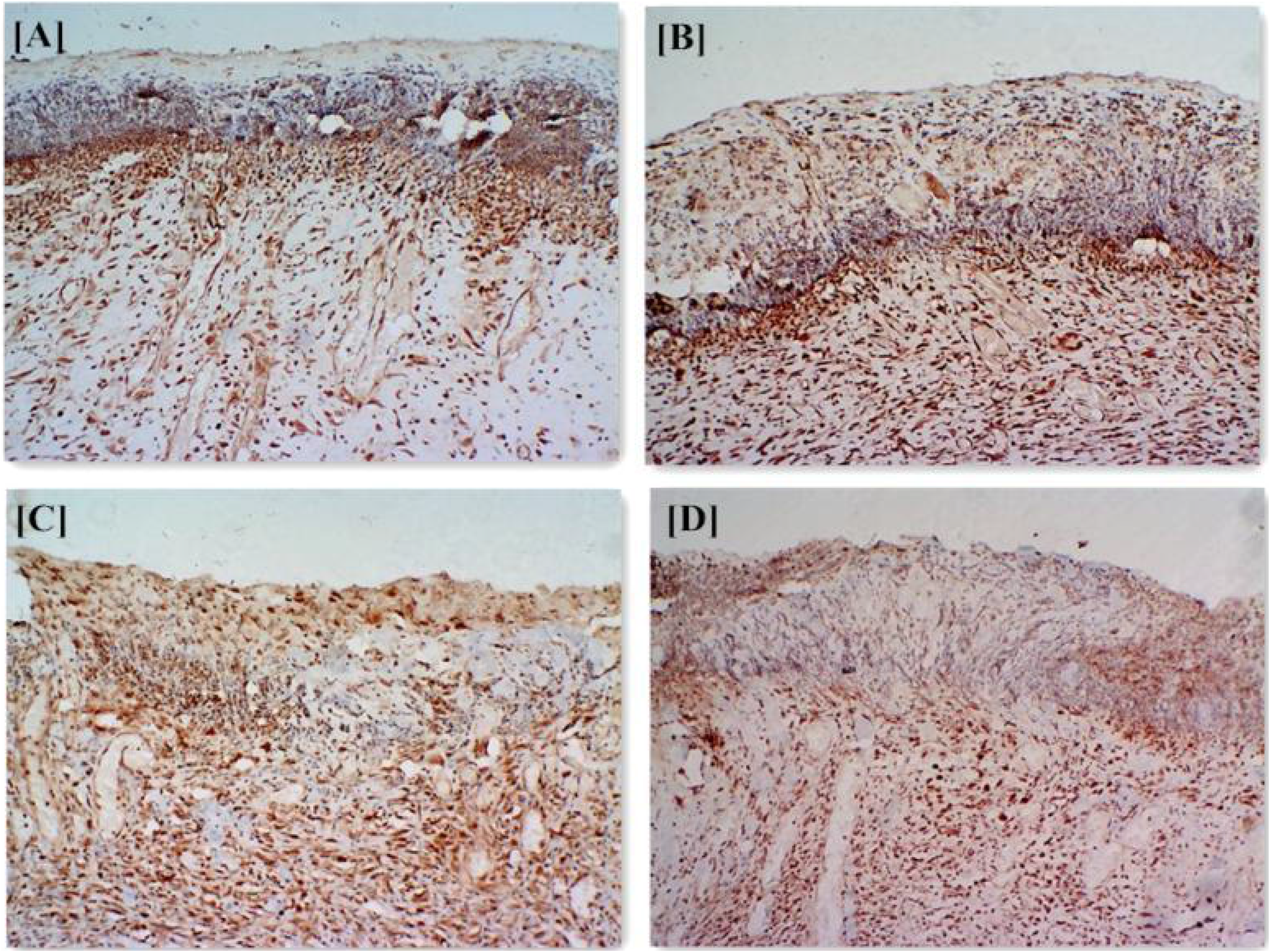

### Comparison of expression levels of TNF-a and IL-1

It is believed that TNF-a and IL-1 are significant inflammatory cytokine which were measured by immunohistochemistry. we used immunoreactivescore to measuring IL-1, which multiply the SI (immune staining)and PP (percentage of positive cells), IPS=SI*PP. SI were graded into four easily reproducible subgroups:(a) no detectable coloration with score of “0”;(b) light yellow coloration with score of “1”;(c)acid coloration with score of “2”,and (d)sepia coloration with score of “3”. PP were graded into 3 groups according to the percentage of positive cell.score of “1” with <30%,score of “2” with 30%-70%,and score of “3” with >70%from the results of IL-1 IHC, It can be see that compared to the control group, treatment groups all showed obvious reduction of IL-1(P<0.05), which means that treatment is beneficial to the reduction of inflammation. in the minocycline combined HA-mediated ultasound group has the best effect compared to the three, the IPS is 3.67(P<0.001),while approach the normal tissue value.the measuring method of TNF-a is count the coloration cells in 400*400 field of vision with randomly 5 field. then count the average number. the results trend is similar to the TNF-a.

## Discussion

Infection wound can cause a series of pathological change in skin tissue[2]. Current therapeutic strategies for hypertrophic scars are not ideal, therefore, basis research in this area has become a hot topic. In this study, we first established a model for infection wound to examine the effects of Minocycline combined hyaluronic acid(HA)-mediated Ultrasound therapy of infected wound in rats. A series of inflammatory factors are positively correlated with the degree of wound infection, such as TNFa, IL-6, IL-1 and etc.and they are sensitive indicators of post-traumatic inflammation, which has the ability to respond to the condition of infection patients, and evaluate the severity of the inflammation[20–21].

Ultrasound can induce structural and functional changes in tissues and cells.according to the the intensity, it can be used to modulate cellular physiological and pathophysiological functions or kill targeted cells [15,16]. Low-level ultrasound sonication has been found to induce reversible changes in membrane permeability and improve drug-delivery efficiency. As such, low-level ultrasound therapy has been used to potentiate the cytotoxicity of chemotherapeutics and improve drug delivery efficiency by sonopermeabilization[22]. Whether used as a sole form of treatment or used combined with other treatments can achieve therapeutic goals. Many studies have confirmed the ability of ultrasound to enhance the effectiveness of chemotherapy drugs[23,24].our lab has previously shown that low-intensity ultrasound has antibacterial effects for killing Staphylococcus aureus [25]. this time we try to combined ultrasound and drugs together, the results showed that there was a significant synergistic effect between ultrasound and drugs.

Minocycline has good antibacterial effect, which can kill the Staphylococcus aureus validly, however, the plasma membrane limits the uptake of minocycline[26], HA is widely used in Clinical Surgery because of ability of skin healing.it can also permeate into the deep tissue somehow. The details of the delivery mechanism have not been studied clearly yet, but it is reasonable to assume a combination of HA binding to cell surface receptors (also for bacteria as seen from the positive effect of HA on their growth) and a pH-induced release of TA[27,28]. As a natural consequence of this mechanistic hypothesis, these particles may hold potential for the treatment of macrophages intracellular pathogens. however, the transport ability is small, it can’t achieve ideal effect. therefore, other mechanisms must play a role.

The ultrasound therapy may accelerate the take-up of drug in two ways. The ultrasound may induce the generation of radicals, which may lead to destabilization of the cell membrane, thereby rendering the cell more susceptible to enhanced drug transport into the cell. another guess is that Minocycline combined hyaluronic acid-mediated ultrasound might relate to sonoporation, drugs can enter the infection cells through induced transient pores, which can reseal following drug entry.

## Conclusions

The goal of our study was to investigate if Minocycline combined hyaluronic acid-mediated Ultrasound has effect in healing infection wound. in vivo testing showed that Minocycline combined hyaluronic acid gel has effect in healing wound, and ultrasound has a little effect in healing wound, combined two for treat infection wound, has great effect conpare with alone. Therefore, there is potential for Minocycline combined hyaluronic acid-mediated Ultrasound to be used in stable, effective, and potentially targetable wound healing measure.

## References

[1] Del Rosso JQ. Wound care in the dermatology office:where are we in 2011? [J]. Am Acad Dermatol 2011;64:1–7.

[2] Li W, Dasgeb B, Phillip T, Li Y, Chen M, Garner W, Woodley DT. Wound-healing perspectives. Dermatol Clin 2005;23:181–92.

[3] Matsukawa A, Yoshimur T, Miyamoto K, et al. Analysis of the inflammatory cytokine network among TNF alpha, IL-1beta IL-1 receptor antagonist, and IL-8 in LPS-induced rabbit arthritis [J]. Lab Invest, 1997, 76(5):629–638.

[4] Eming SA, Krieg T, Davidson JM. Inflammation in wound repair: molecular and cellular mechanisms. [J]. Invest Dermatol 2007;127:514–25.

[5] Werner S, Grose R. Regulation of wound healing by growth factors and cytokines [J]. Physiol Rev, 2003, 83(3):835–870.

[6] Lee JH, Kim HL, Lee MH, You KE, Kwon BJ, et al Asiaticoside enhances normal human skin cell migration, attachment and growth in vitro wound healing model. Phytomedicine, 19(13):1223–7. doi:10.1016/j.phymed.2012.08.002. PMID:22939261.

[7] M. Essendoubi, C. Goblet, R. Reynaud. Human skin penetration of hyaluronic acid of different molecular weights as probed by Raman spectroscopy. [J] Skin Research and Technology, 2016;22:55–62.

[8] Barbucci R, Lamponi S, Borzacchiello A, Ambrosio L, Fini M, Torricelli P, Giardino R. Hyaluronic acid hydrogel in the treatment of osteoarthritis. Biomaterials 2002;23: 4503–4513.

[9] Chen WY, Abatangelo G. Functions of hyaluronan in wound repair. Wound Repair Regen 1999;7: 79–89.

[10] Katoh S, Maeda S, Fukuoka H, et al. A crucial rode of sialidase Neu1 in hyaluronan receptor function of CD44 in T helper type 2-mediated airway inflammation of murine of murine acute asthmatic model [J]. Clinical and experimental immunology, 2010, 161:233–241.

[11] Rafi AQ, Chen D, Schmits R, et al. Evidence for the involvement of CD44 inn endothelial cell injury and induction of vascular leak syndrome by IL-2 [J]. Journal of immunology, 1999,163:1619–1627.

[12] Wang F, Gao Q, Guo S, Cheng J, Sun X, Li Q, Wang T, Zhang Z, Cao W, Tian Y. The sonodynamic effect of curcumin on THP-1 cell-derived macrophages. BioMed Res Int. 2013; 2013:737264.

[13] D.A. Backman, R.L. Brent. Mechanisms of teratogenesis. Ann rev pharmacol Taxicol. 1984, 24:483–500.

[14] Yueping Wang, Deshu Zhuang, Xu Liu et al. Effect of photodynamic therapy on A253 cells of submandibular gland squamous cell carcinoma. [J]. Chinese Journal Practical Stomatology. 2015, 8(12):734–738.

[15] Liu L, Song Y, Ma L, et al. Growth inhibition effect of HMME-mediated PDT on hepatocellular carcinoma HepG2 cells[J]. Lasers Med Sci, 2014, 3(14):14–17.

[16] Salinas GD, Whitworth L, Merwin P, et al. Pancreatic Cancer Management and Treatment Landscape Awareness of Gastoenterologists:Rseults from US Physician Surveys Conducted in 2013-2015.[J]. Gastrointest Cancer:2016, 12(22), doi: 10.1007/s12029-016-9906-5.

[17] Deshu Zhuang, Jialong Han, Liangjia Bi et al. Sonodynamic effect of hematoporphyrin monomethyl ether on ligature-induced periodontitis in rats. [J]. Drug Design, Development and Therapy. 2015:9 2545–2551.

[18] Singer AJ, Homan CS, Church AL, et al. Low-frequency sonophoresis: pathologic and thermal effects in dogs.[J]. Acad Emerg Med. 1998 Jan;5(1):35–40.

[19] Ferrara KW, Borden MA, Zhuang H. Lipid-shelled vehicles: enginerring for ultrasound molecular imaging and drug delivery. Acc Chem Res. 2009 Jul 21;42(7):881–92.

[20] Farwick M, Gauglitz G, Pavicic T, Kohler T, Wegmann M, Schwach Abdellaoui K, Malle B, Tarabin V, Schmitz G, Korting HC. Fifty-kDa hyaluronic acid upregulates some epidermal genes without changing TNF-alpha expression in reconstituted epidermis. Skin Pharmacol Physiol 2011;24: 210–217.

[21] Farwick M, Gauglitz G, Pavicic T, Kohler T, Wegmann M, Schwach Abdellaoui K, Malle B, Tarabin V, Schmitz G, Korting HC. Fifty-kDa hyaluronic acid upregulates some epidermal genes without changing TNF-alpha expression in reconstituted epidermis. Skin Pharmacol Physiol 2011;24: 210–217.

[22] Pack KJ, Mohannad GH et al. A Novel Approach to Ultrasound-Mediated Tissue Decellularization and Intra-Hepatic Cell Delivery in Rats. Ultrasound Med Biol. 2016;42(8):1958–67.

[23] Hasanzadeh H, Mokhtari-Dizaji M et al. Enhancement and control of acoustic cavitation yield by low-level dual frequency sonication: a subharmonic analysis. Ultrason Sonochem. 2011;18(1):394–400.

[24] Cai Y, Fan Y et al. Synergistic effects of ahminoglycosides and fosfomycin on Pseudomonas aeruginosa in vitro and biofilm infection s in a rat model. J Infect Control. 2005;33:78–82.

[25] Zhuang D, Hou C et al. Sonodynamic effects of hematoporphyrin monomethyl ether on Staphylococcus aureus in vitro. FEMS Microbial Left. 2014;361:174–80.

[26] Fabian Nguyen, Agata L. Starosta etc. Tetracycline antibiotics and resistance mechanisms. Biol.chem. 2014;395(5):559–575.

[27] Campo GM, Avenoso A, Campo S, D’Ascola A, Nastasi G, Calatroni A. Molecular size hyaluronan differently modulates toll-like receptor-4 in LPS-induced inflammation in mouse chondrocytes. Biochimie. 2010;92:204–15.

[28] Borzacchiello A Ambrosio L. Network formation of low molecular weight hyaluronic acid derivatives [J]. Biomater Sci Polymer Edn, 2001, 12(3):307–316.

